# RASTRO: Branching Point Decision-Making in Unitig Graphs

**DOI:** 10.64898/2026.09.28.754897

**Authors:** Sana Behtari, Mohsen Hooshmand

## Abstract

**Motivation:** De novo genome assembly relies on accurate path selection through complex graphs, yet unitig graph branching points remain a major ambiguity source. Multiple extensions may be equally plausible from topology alone, but sequence-level consistency of competing paths remains largely unexplored.

**Model:** We introduce RASTRO, a reference-free framework for scoring candidate paths at unitig branching points using nucleotide sequence information. RASTRO-S employs a sample-specific *n*-gram model to estimate extension plausibility via bits-per-base surprisal, while RASTRO-L uses the Evo2 pretrained DNA language model for autoregressive path scoring. Both operate directly on unitig graphs without reference genomes or assembled contigs. To evaluate branch decisions independently, we develop a read-based validation strategy using boundary *k*-mers spanning the junction between graph context and candidate extensions.

**Results:** Experiments on HIV-1 and SARS-CoV-2 unitig graphs from Logan show sequence-based models provide meaningful match with read-supported choices. Among 20,000 sampled branching points, unique read-supported choices were identified for 15,319 HIV-1 branches (76.6%) and 19,882 SARS-CoV-2 branches (99.4%). RASTRO-S improved match from 54.1% to 75.8% for HIV-1 and from 58.6% to 76.4% for SARS-CoV-2 as context length rose from *n* = 5 to *n* = 10. RASTRO-L also improved with context length, reaching 49.96% and 49.25% match. Both outperformed random selection and the longest-extension greedy baseline.

**Conclusion:** RASTRO provides an efficient, interpretable framework for evaluating ambiguous path decisions in unitig graphs using raw-read evidence rather than contig-level alignment. Results demonstrate the potential of lightweight, sample-adaptive sequence models for improving local decision-making in genome assembly workflows.

## Introduction

Advances in next-generation sequencing (NGS) technologies have dramatically increased the volume of genomic data while reducing sequencing costs, making whole-genome sequencing widely accessible across biological and medical research (Ang et al., 2025). In *de novo* genome assembly, the objective is to reconstruct an organism’s genome directly from sequencing reads without relying on a reference genome (Sohn and Nam, 2018). Accurate *de novo* assembly is essential for studying previously uncharacterized organisms, identifying structural variations, and analyzing highly diverse microbial and viral populations. Despite substantial progress in sequencing technologies and assembly algorithms, reconstructing complete genomes from short sequencing reads remains a challenging computational problem (Thomma et al., 2016). Most modern *de novo* assemblers address this challenge by constructing a de Bruijn graph, in which edges represent *k*-mers extracted from sequencing reads and nodes indicate consecutive (*k* + 1)-mer overlaps between them (Pevzner et al., 2001) (Compeau et al., 2011). To simplify the graph and reduce its complexity, maximal non-branching paths are compressed into unitigs, resulting in a compact graph representation while preserving its connectivity. Several methods have specifically addressed the efficient construction and compaction of these unitig graphs. BCALM2 (Chikhi et al., 2016) partitions *k*-mers using left and right minimizers, compacts the resulting buckets independently, and then reunites the compacted sequences to obtain maximal unitigs. Cuttlefish (Khan et al., 2025) instead models de Bruijn-graph vertices as finite-state automata and uses implicit graph traversals to determine vertex states and extract maximal unitigs without explicitly constructing the full graph. Assemblers such as SPAdes (Bankevich et al., 2012) further construct and simplify assembly graphs using multisized de Bruijn graphs, together with procedures for removing or correcting graph structures arising from sequencing errors and variation. Despite the substantial progress in unitig construction and graph simplification, branching points remain unresolved. Graph construction tools mostly terminate unitig extension whenever multiple outgoing paths are encountered, preserving these ambiguities rather than attempting to resolve them. Despite the simplification done by unitigs, branching points remain unresolved. Graph construction tools mostly terminate unitig extension whenever multiple outgoing paths are encountered, preserving these ambiguities rather than attempting to resolve them. A decision point refers to a unitig strand from which at least two distinct outgoing edges originate. Such points represent branching locations in the unitig graph where the available graph structure alone does not determine a unique continuation. Resolving a decision point therefore requires selecting one of the alternative outgoing paths as the most appropriate continuation of the current unitig strand. The presence of multiple alternatives can arise from repetitive sequences, sequencing errors, genetic variation, or other sources of ambiguity in the underlying sequencing data. Consequently, decision points constitute critical locations for resolving graph-level ambiguities during genome assembly. (Chikhi et al., 2016) (Khan et al., 2025). Downstream assemblers subsequently select among the competing continuations using heuristics such as paired-end information, read coverage, or graph topology (Bankevich et al., 2012). While these strategies have proven effective in many cases, branch selection remains one of the most challenging steps in genome assembly, particularly in repetitive or highly variable genomic regions where multiple alternatives may appear equally plausible (Salmela, 2025).

Statistical approaches have also been explored to incorporate uncertainty and probabilistic evidence into genome assembly. By modeling the likelihood of sequencing data or the probability of alternative assembly configurations, these methods can provide a principled basis for evaluating competing solutions beyond graph topology alone (Boža et al., 2015) (Howison et al., 2014). Although such approaches demonstrate the potential of statistical modeling for handling ambiguity in genome assembly, their application to sequence-level decision-making at individual branching points within unitig graphs has received comparatively limited attention.

In recent years, the intersection of artificial intelligence and bioinformatics has led to a transformative paradigm: treating biological sequences as a form of language (Veiner and Supek, 2026) (Feng et al., 2025). Just as words follow grammatical rules to form meaningful sentences in natural languages, nucleotide sequences adhere to complex biological constraints and sequential patterns (Zhou et al., 2024) (Dalla-Torre et al., 2025) (Brixi et al., 2026). While advanced statistical and deep learning-based language models have been increasingly adopted for tasks such as functional annotation, regulatory region identification, and sequence classification, their potential for resolving local structural ambiguities, specifically in guiding path selection at branching points within assembly graphs, remains largely unexplored.

To address this gap, we introduce RASTRO, a framework comprising two sequence-based methods. The first, RASTRO-*S*, is a sample-driven *n*-gram statistical model that learns local sequence patterns from the unitigs of the target sample and assigns information-theoretic scores, expressed in bits per base (*bpb*), to candidate extensions. These scores are derived from conditional probabilities, with lower values indicating greater consistency with the patterns captured by the model. The second, RASTRO-*L*, extends this framework using the pre-trained genomic language model Evo2 (Brixi et al., 2026) to assess the plausibility of competing extensions based on learned genomic sequence representations.

Together, the two methods provide sequence-based criteria for local branch selection without requiring a reference genome or a completed assembly. Both are tunable, allowing the decision process to be adjusted according to the desired level of selectivity, while their computational requirements depend primarily on the input sequence length and the number of branching points in the graph.

The main contributions of this work are:

- a sample-specific *n*-gram framework for quantifying the sequence-level plausibility of competing extensions at unitig graph branching points;
- an LLM-based approach of this framework using the pre-trained genomic language model Evo2 (Brixi et al., 2026);
- a read-based evaluation framework for assessing branch-selection decisions directly against sequencing evidence, together with an analysis of the effects of sequence context and evidence strength.

This paper is structured as follows. Section 2 reviews prior work on branching-point resolution in assembly graphs. Section 3 presents the Methods, including the viral datasets obtained from Logan and the two proposed branch-scoring approaches, RASTRO-S and RASTRO-L. Section 4 reports the experimental results, covering evaluability rates and match with read-supported choices. Finally, Section 5 concludes the paper and outlines directions for future research on integrating sequence-driven branch scoring into assembly workflows.

## Related work

Branching points in de Bruijn and unitig graphs represent one of the primary sources of ambiguity in genome assembly, as multiple graph continuations are equally consistent with the graph topology alone. Consequently, graph construction tools such as BCALM (Chikhi et al., 2016) and Cuttlefish (Khan et al., 2025) construct unitigs by extending maximal non-branching paths and terminating the extension when a branching point is encountered. Rather than resolving these ambiguities, they preserve them for downstream assembly stages.

Several assemblers subsequently attempt to resolve these branching points using additional sequencing evidence. Velvet (Zerbino and Birney, 2008), ABySS (Simpson et al., 2009), and SPAdes (Bankevich et al., 2012) all begin by simplifying the assembly graph through the removal of sequencing errors such as tips, bubbles, and low-coverage branches before constructing unitigs from unambiguous paths. The remaining ambiguities are then handled primarily through paired-end information. In all three approaches, branch resolution depends on external read-pair evidence rather than directly evaluating the biological plausibility of competing sequence continuations.

Other methods formulate branch resolution as a probabilistic optimization problem. GAML (Boža et al., 2015) searches for the assembly that maximizes the likelihood of the observed sequencing data using simulated annealing, allowing alternative graph traversals to compete during optimization while avoiding unsupported decisions when multiple solutions remain equally likely. GABI (Howison et al., 2014) extends this idea by treating genome assembly as Bayesian inference, using Markov Chain Monte Carlo (MCMC) sampling to generate a posterior distribution over alternative assemblies rather than producing a single deterministic result. Similarly, SAMA (Salmela, 2025) estimates the probability of misassembly for graph edges and constructs contigs only when the estimated risk remains below a user-defined threshold. These approaches explicitly model uncertainty during assembly but operate at the level of complete assemblies or graph traversals rather than evaluating individual branch continuations from sequence context.

Recent advances in genomic foundation models provide a complementary source of information for resolving branching points. Instead of relying on graph topology or paired-end connectivity, DNA language models learn statistical regularities directly from genomic sequences and estimate the likelihood of candidate continuations (Veiner and Supek, 2026) (Feng et al., 2025). DNABERT-2 (Zhou et al., 2024) introduced an efficient genome language model based on Byte Pair Encoding (BPE), substantially reducing computational cost while achieving strong performance across diverse genome understanding tasks. Nucleotide Transformer (NT) (Dalla-Torre et al., 2025) further developed this paradigm through a family of pretrained models ranging from millions to billions of parameters, trained on large collections of genomic sequences and evaluated across a broad range of genomic prediction tasks. More recently, Evo2 (Brixi et al., 2026) scaled genomic language modeling to trillions of nucleotides spanning all domains of life, demonstrating the ability of large-scale models to capture biological sequence patterns across diverse organisms and tasks. Despite these advances, genomic language models have primarily been evaluated on sequence understanding, functional prediction, and variant analysis, and their potential for guiding branch selection during genome assembly remains largely unexplored.

Existing evaluation methodologies likewise focus predominantly on the quality of completed assemblies or corrected sequencing reads. Metrics such as contiguity, misassembly rate, and consensus accuracy assess the final assembly, while tools such as LRCstats (La et al., 2017) evaluate read-correction accuracy by comparing corrected reads against reference sequences. Although these approaches effectively measure end-to-end assembly quality, they do not directly evaluate whether a branch-selection strategy chooses the continuation most strongly supported by the underlying sequencing reads. This motivates branch-level evaluation, where competing continuations are assessed independently of the final assembly and compared directly against evidence contained in the raw sequencing reads.

## Methods and material

This section describes the datasets, graph representations, and methodological procedures used to develop and evaluate RASTRO for branch decision-making in unitig graphs. We first describe the genomic datasets and corresponding unitig graphs and raw sequencing reads used in our experiments. We then present the two proposed branch-scoring approaches, RASTRO-S and RASTRO-L, finally followed by the read-based evaluation procedure used to assess their decisions against sequencing evidence.

### Dataset

We sourced our data from Logan (Chikhi et al., 2024), a genomic repository containing pre-constructed unitig graphs alongside their corresponding raw sequencing reads. Utilizing Logan significantly simplifies our methodology by eliminating the computationally intensive need to construct assembly graphs from scratch. For our evaluation, we selected the HIV-1 and SARS-CoV-2 viral datasets extracted from this repository, selected because of their available raw sequencing reads and suitability for unitig-graph construction and read-based branch evaluation (Dingens et al., 2019; Baker et al., 2021). Figure 1 illustrates an excerpt from the Logan dataset, showing a unitig sequence file (SRR7693968.unitig.fa) used in our experiments. Each entry corresponds to a unitig record containing sequence identifiers, positional metadata, and nucleotide content, which serve as input for constructing candidate extensions in the RASTRO framework.

**Figure 1.**
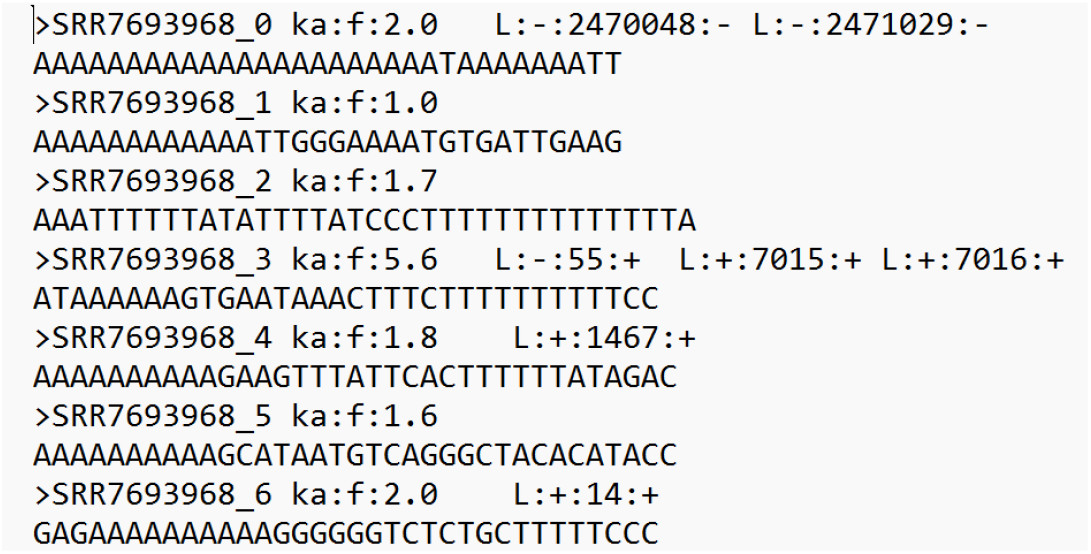
Example unitig record from the Logan dataset (SRR7693968.unitig.fa).

### RASTRO-S: sample-specific n-gram scoring model

The primary objective of RASTRO-*S* is to provide a lightweight and fast scoring mechanism for path selection at branching points in a unitig graph, as illustrated in Figure 2. To capture the local sequence statistics of each dataset, RASTRO-*S* is trained on all unitigs contained in the corresponding Logan FASTA file. For every candidate extension at a branching point, the model computes the entropy per base (bits-per-base), which is used to rank competing paths. Lower bits-per-base values indicate greater statistical consistency with the sequence composition of the dataset.

**Figure 2.**
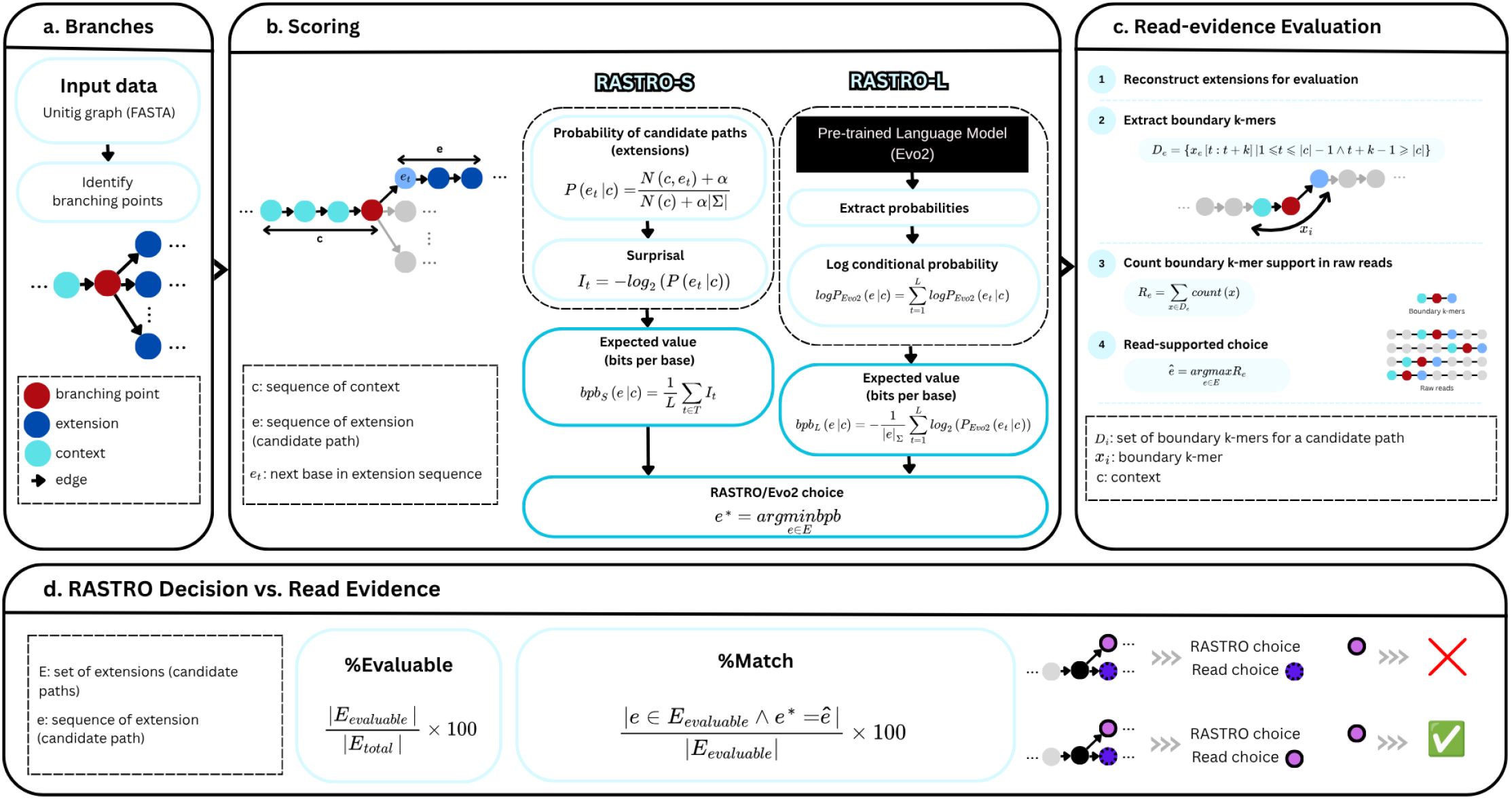
Overview of RASTRO framework for branching-point decision-making in unitig graphs. (a) Branches: Branching points are identified from the unitig graph obtained from Logan. (b) Scoring: Candidate extensions are scored by RASTRO-S (sample-specific n-gram model) and RASTRO-L (Evo2), both producing bits-per-base (*bpb*) scores; the extension with the lowest *bpb* is selected. (c) Read-evidence Evaluation: Boundary k-mers spanning the context–extension junction are extracted and counted in the raw sequencing reads to determine the read-supported choice. (d) RASTRO Decision vs. Read Evidence: Model decisions are compared with the read-supported choice to compute the percentage of evaluable branches and the match rate.

The model’s alphabet is defined as Σ = {*A, C, G, T*}. Any character outside of this set is excluded from both training and scoring. RASTRO-*S* is an *n*-th order Markov model, where the probability of the next base depends solely on the *n* preceding bases. For every context **c** ∈ Σ^*n*^ of length *n* and the subsequent base *e*_*t*_ ∈ Σ, we estimate the following conditional distribution:

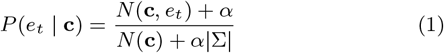

where *N* (**c**, *e*_*t*_) is the number of occurrences of the context **c** with length *n* immediately followed by base *e*_*t*_ across the entire unitig, 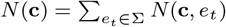 is the total number of occurrences of context **c**, *α* = 1 is the Laplace smoothing parameter used to prevent zero probabilities, |Σ| = 4.

If *N* (**c**) = 0, meaning that the context **c** has not been observed in the training corpus, the conditional probability is estimated from the unigram distribution instead of assigning a uniform probability:

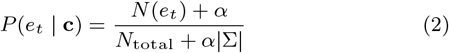

where *N* (*e*_*t*_) is the number of occurrences of base *e*_*t*_ in the corpus, and 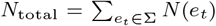 is the total number of observed bases. This fallback strategy allows the model to rely on the overall nucleotide composition of the sample when contextual information is unavailable.

If *S* is a training sequence of length |*S*| (after excluding characters outside of Σ), each valid position *i* defines a training example consisting of an *n*-base context and the subsequent base. Specifically, for

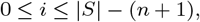

the context and subsequent base are defined as

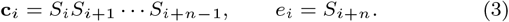

Thus, **c**_*i*_ contains the *n* bases preceding *e*_*i*_, while *e*_*i*_ is the next base to be predicted. Each context–base pair is then used to update the corresponding transition count:

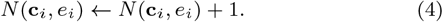

These counts are aggregated across all unitigs and subsequently converted into conditional probabilities using the equations defined above. In addition, when a context **c** has not been observed in the training corpus, the overall base counts *N* (*e*_*t*_) and their total *N*_total_ are obtained from the same corpus and used for the unigram fallback distribution.

As explained in Section 1, a decision point is defined as a unitig strand, consisting of a unitig *u* and strand orientation *σ* ∈ {+, −}, with at least two outgoing edges, Figure 2, a. For each decision point, the suffix of the source unitig in the direction *σ* is extracted and provided to the model as the context (prefix). A candidate extension is then generated for each outgoing edge by traversing the graph forward along linear paths. The traversal terminates when another branching point is encountered or when predefined traversal limits are reached. Each context–extension pair is subsequently scored, and the resulting scores are used to rank the competing candidate paths.

Consequently, for each branch, we obtain a set of candidate options, each representing a probable continuation from that branch. To score base *e*_*t*_, we define **c** as the *n* bases immediately preceding it, as mentioned before. This definition is applicable when the length of the string formed prior to *t* is at least *n* bases. Otherwise, scoring initiates from the first base that fulfills this condition. Next, the probability of the subsequent base is obtained from the model in Equation 1. The probability assigned by the model can be converted into an information-theoretic quantity known as surprisal (Hale, 2016). Surprisal measures the information content of an observed event and is given by the negative logarithm of its probability:

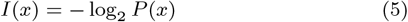

Thus, lower-probability events have higher surprisal values. In the context of RASTRO-*S*, the event of interest is the observation of a specific nucleotide *e*_*t*_ given its preceding context **c**. Therefore, specializing the general definition in Equation 5 to the conditional probability assigned to *e*_*t*_ gives the position-specific surprisal:

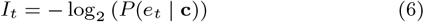

The final score for a candidate extension is computed as the average surprisal across all scorable positions:

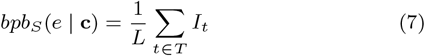

where *T* is the set of indices for which a context of length *n* can be formed, and *L* = |*T* |. Lower *bpb*_*S*_ values indicate that the extension is more consistent with the sample-specific sequence statistics learned by the model and are therefore ranked more favorably.

As mentioned above, RASTRO-*S* is based on an *n*-th order Markov chain that models the dependency of each nucleotide on the preceding *n* bases, Algorithm 1. During training, all unitigs in the dataset are traversed once to collect the frequencies of observed *n*-grams and their transitions. If the total length of the training corpus is *N*, the training time complexity is therefore *O*(*N*).

For a fixed model order *n*, the state space consists of at most 4^*n*^ contexts and 4^*n*+1^ transitions. Consequently, the memory requirement depends only on the model order and the DNA alphabet, rather than the genome size, making RASTRO-*S* computationally efficient in practice.

In the scoring phase, the probability of each base is retrieved from a pre-computed lookup table, requiring only a linear traversal of the candidate sequence. Consequently, scoring an extension of length *L* has a time complexity of *O*(*L*). If a branching point has *E* candidate extensions, the cost of evaluating that branching point is *O*(*EL*). Assuming the graph contains *M* branching points, the overall scoring complexity is therefore *O*(*MEL*).

Since RASTRO-*S* relies solely on table lookups and simple arithmetic operations, its computational overhead is substantially lower than that of large language models, making it well suited for large-scale branch evaluation on standard computing hardware.

### RASTRO-L: Evo2-based branch scoring

Large language models (LLMs) have demonstrated the ability to capture complex sequential dependencies by learning statistical patterns from large-scale sequence data (Feng et al., 2025). In genomics, pretrained DNA language models extend this principle to nucleotide sequences, providing the means of estimating the likelihood of candidate bases or sequence continuations based on their preceding context (Dalla-Torre et al., 2025) (Brixi et al., 2026). This makes autoregressive genomic language models particularly relevant to branch selection, where competing extensions can be evaluated according to how consistent they are with learned sequence patterns rather than graph structure alone.

As illustrated in Figure 2, RASTRO-*L* utilizes Evo2 to score the candidate paths, which is an auto-regressive model trained with the objective of next-token prediction (similar to text language models, but with the distinction that the units are DNA bases). The output of the model at each step is “next-token prediction.”

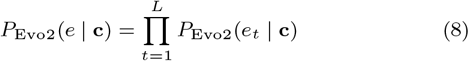

where *P* (*e*_*t*_ | **c**) denotes the next-token probability produced by the model at step *t*. In other words, by receiving context, it produces a probability distribution over {*A, C, G, T*} for the next base. In this study, the output is not used for “sequence generation,” but rather for scoring a specific extension *e* such that, given context **c** and extension *e* at step *t*, the model observes the current context and generates the probability of the next base. Then, among these probabilities, only the value corresponding to the actual extension base is retrieved. By aggregating these values over the length of the extension, the conditional log-probability of the entire extension is obtained, followed by its normalized version.

#### Algorithm 1

RASTRO-*S*: Sample-Specific *n*-gram Scoring

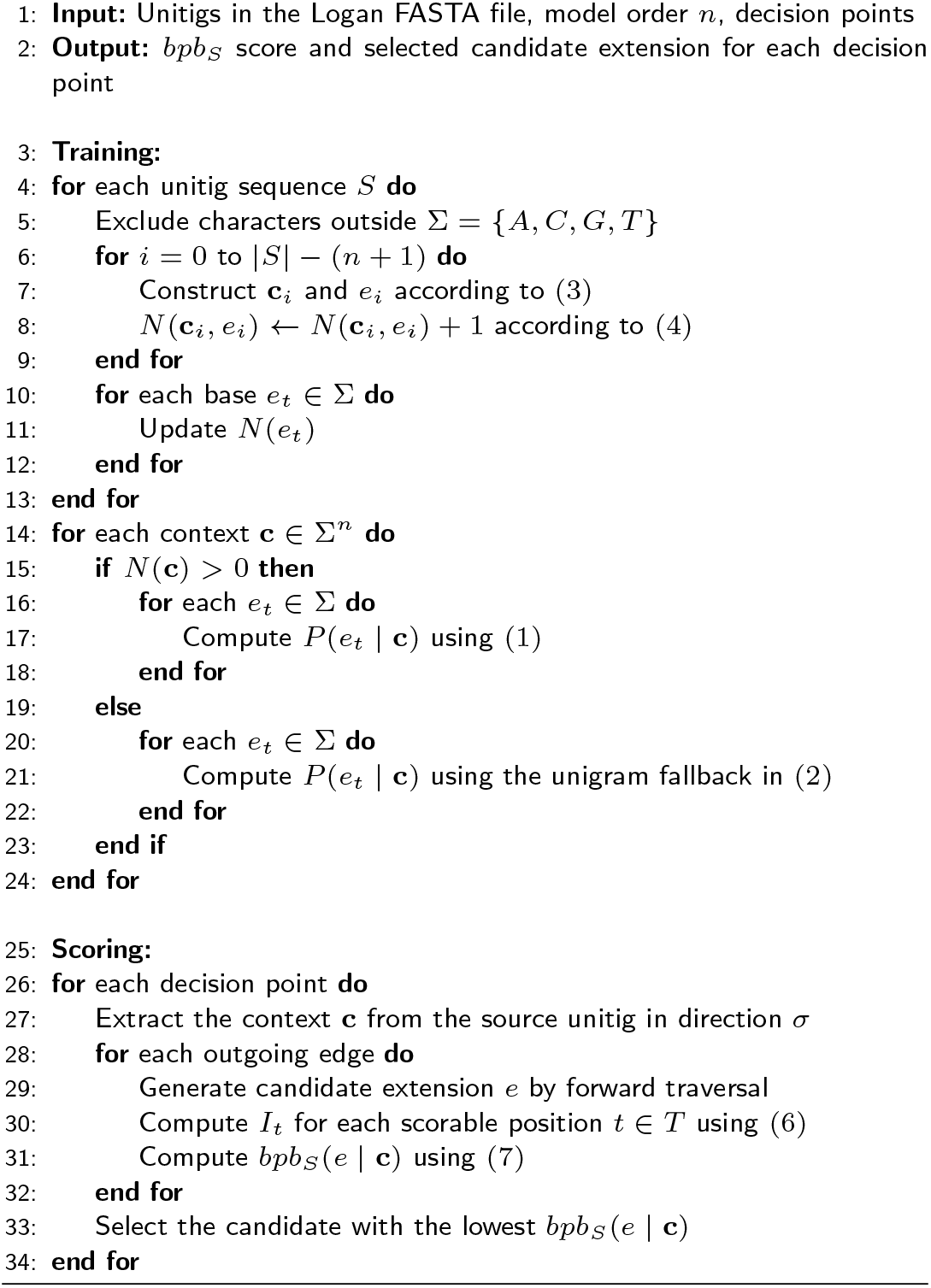

Evo2 autoregressively computes the probability of each extension base conditioned on context bases. The log conditional probability of the entire extension is defined as follows:

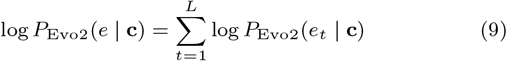

At step *t*, the model has access to a sequence formed by context and extension bases up to that point.

To ensure that candidates of varying lengths can be fairly compared, the scores obtained from the previous equation are normalized by length. That is, in addition to storing log-probability values, the *bpb* metric is also calculated.

And let |*e*|_Σ_ be the number of extension bases that belong to Σ. Then:

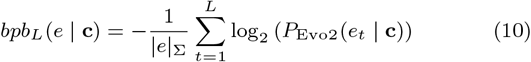

In this definition, a smaller value indicates a more probable (and consequently more natural) extension, Algorithm 2.

#### Algorithm 2

RASTRO-*L*: Evo2-Based Branch Scoring

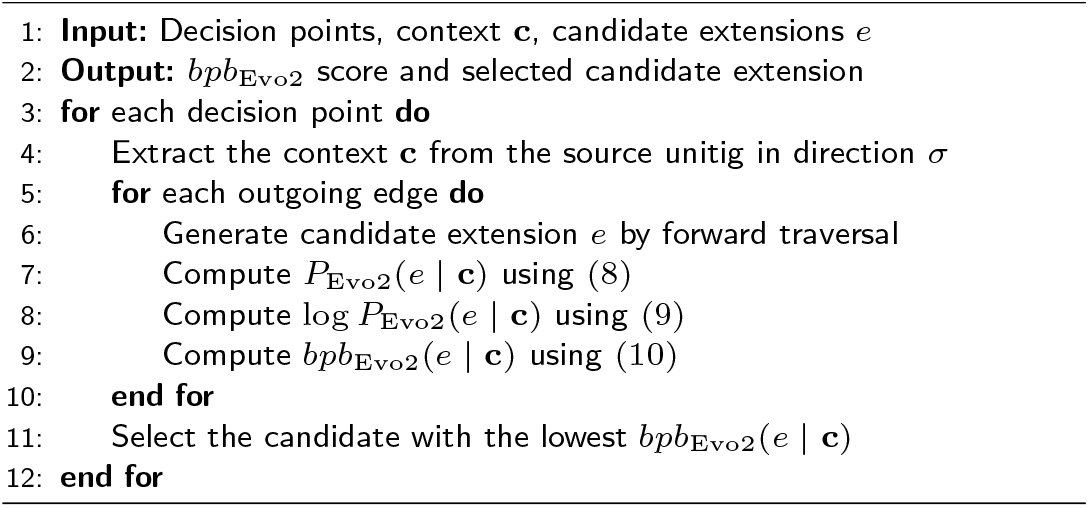

### Read-based Evaluation

Based on the branch scoring table, the Logan unitig graph, and the raw sequencing reads, the evaluation procedure consists of the following steps: (1) loading the unitig graph and constructing the minimal traversal structures required for candidate-path exploration; (2) identifying branching points and generating all valid candidate extensions; (3) selecting the candidate with the lowest *bpb* score whenever a branch contains at least two comparable alternatives; (4) determining the read-supported choice by extracting boundary *k*-mers for each candidate path and counting their occurrences in the raw sequencing reads; and (5) evaluating the model by reporting the percentage of scorable branches and the match between model decisions and read-supported choices under different confidence thresholds.

We evaluate branch selection independently of the final assembled contigs by using the original sequencing reads as the reference source of evidence. Reads are the direct output of the sequencing process and therefore remain independent of the assembly algorithms and heuristics used for graph traversal. In contrast, contigs are themselves the result of branch selection decisions and may therefore obscure ambiguities or introduce algorithm-specific bias. Using reads as the evaluation reference provides an independent measure of branch-selection quality based on direct experimental evidence from the sample.

For each branching point, multiple candidate extensions may exist. Since a lower *bpb* score indicates a more likely extension, the model selects the candidate with the minimum score:

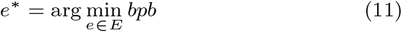

where *E* denotes the set of candidate extensions for branch *e*.

If two or more candidate extensions receive identical scores, the tie is broken deterministically by selecting the lexicographically smaller extension.

Only branches for which at least two candidate extensions receive scores are included in the evaluation. The percentage of evaluable branches is computed as

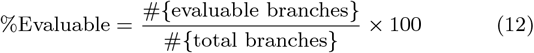

*Evaluable rate*, represents the percentage of branches for which a valid model decision can be obtained. To obtain an independent reference for evaluation, a *read-supported choice* is determined for each evaluable branch. A short suffix of the source unitig is first extracted as the context. This context is concatenated with each candidate extension to form a unique query sequence representing that branch transition. Boundary *k*-mers are then generated from each query sequence, and their occurrences are counted in the raw sequencing reads. The candidate whose boundary *k*-mers receive the highest support is selected as the read-supported choice and serves as the reference for evaluating the model’s decision.

Since the objective is to measure support for the transition from the context to a candidate extension, rather than for the extension sequence alone, only boundary *k*-mers are considered. These *k*-mers overlap both the suffix of the context and the prefix of the extension, thereby uniquely representing the junction between them. Consequently, matches in the raw sequencing reads provide direct evidence for the corresponding branch transition while reducing spurious support from identical sequences occurring in different contexts.

By having **c** as the context and *e* as the extension corresponding to the specific candidate, we first construct the concatenated sequence

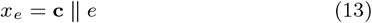

where ∥ denotes string concatenation. The set of boundary *k*-mers for candidate *e* is then defined as

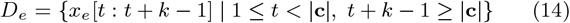

That is, each boundary *k*-mer overlaps both the suffix of the context and the prefix of the extension.

After constructing the set *D*_*e*_ for each candidate extension, containing the *k*-mers that span the junction between the context and the extension, we measure their support in the raw sequencing reads. First, the frequencies of all *k*-mers are computed over a fixed subset of reads. For each boundary *k*-mer, its occurrence count is then retrieved from frequency table.

For each candidate *e*, the read support score is defined as the sum of the occurrence counts of its boundary *k*-mers:

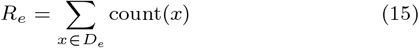

where count(*x*) denotes the number of occurrences of *x* in the read-derived frequency table. Candidates whose boundary *k*-mers receive higher support from the reads therefore obtain larger read scores.

The read-supported choice for each branch is then defined as the candidate with the maximum read score:

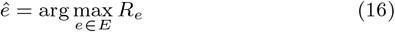

To avoid unreliable selections and analyze the effect of evidence strength, results are additionally reported across different confidence thresholds. These thresholds are based on the ratio between the highest read score and the second-highest read score. Larger ratios indicate stronger separation between candidate extensions and therefore stronger read support.

After obtaining (1) the model decision from the scoring table and (2) the read-supported choice from the sequencing reads for each branch, the match between the model and the read evidence is evaluated. Since the objective is to measure decision quality only on branches where the read evidence provides a clear reference, match is computed only for branches with a unique read-supported choice and a valid model decision (i.e., branches containing at least two candidate extensions with valid and comparable scores).

For a branch *e*, let *e*^*^ denote the model decision and *ê* denote the read-supported choice. The match of the model decision for branch *e* is defined as a binary variable:

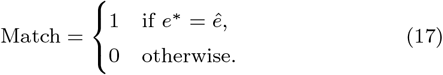

The overall match ratio is then calculated as the fraction of evaluated branches where the model decision matches the read-supported choice:

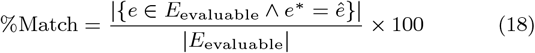

*Match rate*, measures the consistency between the model decision and the read-supported choice among branches where both a valid model decision and a unique read-supported choice are available.

Proposed metrics provide a systematic assessment of branch-selection decisions based on their match with sequencing-read evidence, Algorithm 3. High match rates indicate branches where the model selection is strongly supported by the reads, suggesting more reliable decisions. Conversely, low match rates highlight branches where the model decision is not supported by the available read evidence, identifying potential ambiguous regions or assembly errors that may require further investigation.

#### Algorithm 3

Read-Based Evaluation of Branch Decisions

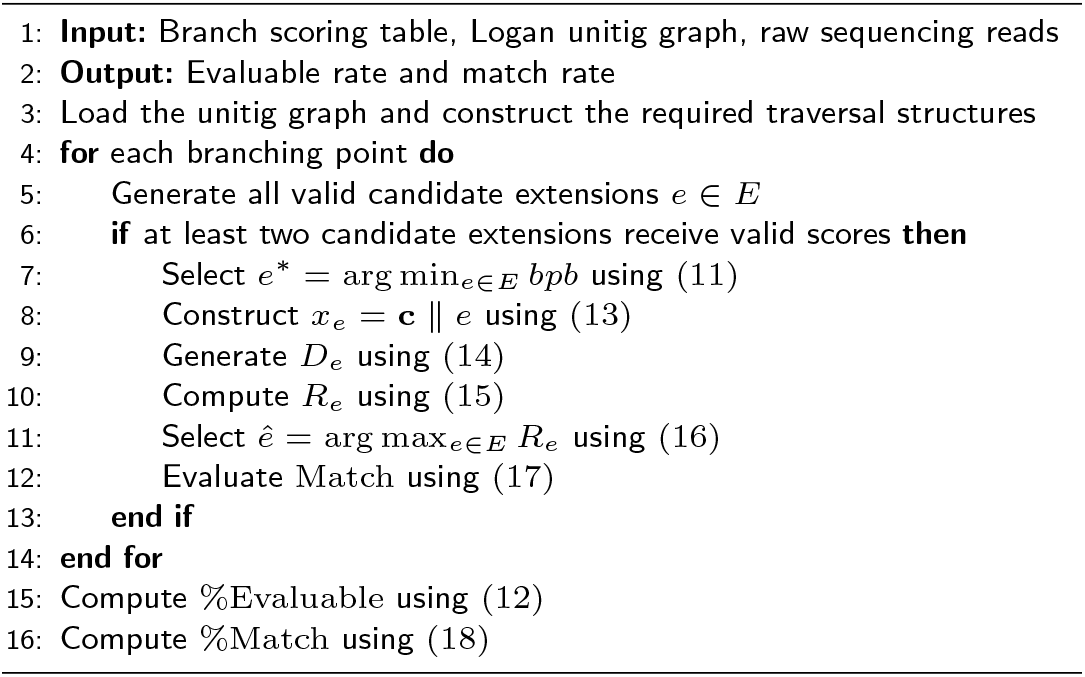

## Results

Most experiments were conducted in the Google Colab Pro environment. The computationally lightweight components of the framework, including candidate-extension extraction from the unitig graph, RASTRO-*S* training and scoring, and the read-based evaluation using *k*-mer counts generated with Jellyfish (Marçais and Kingsford, 2011), were executed on CPU resources. In contrast, RASTRO-*L* scoring requires substantially greater computational resources because of the size and computational cost of the pretrained model (`evo2_7b_base`). In this experiments, NVIDIA A100-class GPUs and the NVIDIA RTX PRO 6000 provided a suitable environment for Evo2, particularly in terms of GPU memory and execution stability compared with lower-memory options such as the T4. However, GPU availability in Colab is variable, and execution may be affected by GPU-memory limitations, session resets, and runtime limits. These constraints have a substantially smaller impact on RASTRO-*S* and the read-based evaluation, which require comparatively limited computational resources.

To evaluate the proposed method, two samples were used: `SRR7693968`, corresponding to human immunodeficiency virus type 1 (HIV-1), and `SRR10605873`, corresponding to SARS-CoV-2. The required graph data, including unitigs and their links, were obtained from the Logan dataset for the corresponding samples. Rather than constructing the unitig graphs directly from the raw reads, the precomputed Logan outputs were used to access the unitig-graph structure and its branching points directly. Subsequently, the raw reads corresponding to the same samples were used independently to determine the read-based choice at each branching point and to compare it with the decisions produced by the scoring models. This read-based evaluation provides evidence that is independent of the final assembly and does not require a reference genome.

The selection of the HIV-1 sample was motivated by several considerations. First, its relatively small genome makes the execution of the proposed method, including candidate-extension generation, scoring, and evaluation, more computationally manageable. Second, the availability of high-quality raw sequencing reads provides the data required to construct the read-based evidence used in the evaluation. Finally, the substantial sequence diversity observed within HIV-1 samples can lead to increased local ambiguity and branching in the unitig graph, making it a suitable dataset for evaluating branch-level decision-making (Dingens et al., 2019). To broaden the evaluation and examine the consistency of the results across different datasets, a SARS-CoV-2 sample was also included. SARS-CoV-2 provides a biologically distinct dataset from HIV-1 and allows the proposed method to be evaluated on a second viral genome with different sequence characteristics. The availability of sequencing data for SARS-CoV-2 also facilitates the application of the same graph-based and read-based evaluation procedure. Including this sample therefore provides an additional basis for assessing whether the observed behavior of the scoring models is consistent across datasets rather than being specific to the HIV-1 experiment (Baker et al., 2021).

For each branch, the candidate extensions are scored independently by RASTRO-*S* and RASTRO-*L* using corresponding *bpb*_*S*_ and *bpb*_*L*_. As mentioned in Section 3, a lower score indicates a higher model-assigned probability and therefore a more favorable candidate under the corresponding model. Table 1 shows an example of the branch-scoring output at a branching point in `SRR7693968`, HIV-1, there exist two branches `SRR7693968_1413392` and `SRR7693968_1413394`. RASTRO-*S* and RASTRO-*L* both correctly select `SRR7693968_1413394`, the same extension, since it receives the lower score under both models.

**Table 1.** Example of branch-scoring output. Two candidate extensions are scored using RASTRO-*S* and RASTRO-*L*. For each model, the candidate with the lower *bpb* score is selected.

| Branch | Extension | RASTRO-*S* | RASTRO-*L* |
| --- | --- | --- | --- |
| SRR7693968.1403155 | SRR7693968.1413392 | 3.328 | 1.702 |
|  | SRR7693968.1413394 | **1.703** | **1.570** |

To evaluate the performance of the proposed method within an interpretable framework, two baseline methods were implemented. The first baseline is a random selection strategy, which estimates the chance-level performance of path selection at branching points. The second baseline is a longest-path greedy strategy, which evaluates whether a simple structural heuristic based on extension length can provide useful branch decisions. These baselines use the same branches, candidate extensions, and read-based evaluation procedure as the proposed method, while differing only in their decision mechanisms. The random baseline is described first, followed by the greedy baseline.

The random baseline was defined to estimate the chance level of path selection at branching points. The baseline uses the same branches, candidate extensions, and read-based evaluation procedure as the proposed method, except for the decision mechanism. For each branch with at least two valid candidate extensions, one candidate is selected uniformly at random. Comparing model match with the random baseline indicates whether the model-based decisions provide information beyond chance-level selection. The random baseline was evaluated using the same experimental setup as the model-based experiments.

A fixed set of 20,000 sampled branches and their corresponding candidate extensions was used. For each branch with at least two valid candidates, one candidate was selected uniformly at random. A fixed random seed was used for the initial experiment to ensure reproducibility. For the HIV-1 sample, 15,319 branches were scorable for comparison with the read-based choice. The random baseline agreed with the read-based choice in 6,127 cases, corresponding to an match rate of approximately 40%. For the SARS-CoV-2 sample, 19,882 branches were scorable, and the random baseline agreed with the read-based choice in 8,556 cases, corresponding to an match rate of approximately 43%. To assess the stability of the random baseline and reduce dependence on a particular random realization, the experiment was repeated for 100 independent runs without fixing the random seed. The mean match was 0.397 ± 0.004 for HIV-1 and 0.427 ± 0.003 for SARS-CoV-2. These values are consistent with the corresponding single-run results obtained using the fixed seed, indicating that the reported baseline is not strongly dependent on a particular random selection. To further validate the obtained baseline values, the expected match under random selection was calculated independently. Since the number of candidate extensions varies across branches, the probability of selecting the read-supported candidate by chance for a branch with *m* candidate extensions is 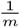. Therefore, the overall expected random match was calculated by averaging 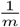 across the set of comparable branches. This calculation resulted in an expected match of 0.396. Its close correspondence with the observed random-baseline match values of approximately 0.40 for HIV-1 and 0.43 for SARS-CoV-2 confirms that the implemented random baseline behaves consistently with the expected statistical chance level.

We also introduce a longest-path greedy strategy for a better comparison. At each branching point, the method considers the same set of candidate extensions generated for RASTRO-*S* and RASTRO-*L* and selects the extension with the greatest sequence length:

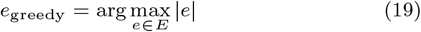

The intuition is that, in the absence of any sequence-statistical or read-based evidence, a locally longer continuation may provide a simple heuristic for selecting between competing paths. The method therefore relies solely on the structure of the locally generated candidate extensions and does not use raw sequencing reads or a language model. If multiple candidates have the same maximum length, the lexicographically smallest extension is selected to ensure deterministic behavior. In the evaluation framework, this heuristic achieved match rates of 24.7% for the HIV-1 dataset and 32.8% for the SARS-CoV-2 dataset, indicating that extension length alone provides limited information for identifying read-supported branch decisions.

Read support is computed for all sampled branches; however, not all branches yield a unique read-supported selection. This is an inherent consequence of the evaluation procedure and the ambiguity of the underlying sequencing evidence, rather than an implementation limitation. Read support is determined using boundary *k*-mers, i.e., substrings spanning the junction between the branch sequence and the beginning of a candidate extension. This restriction ensures that the evidence specifically reflects support for connecting a particular branch to a particular extension, rather than the independent abundance of the extension elsewhere in the dataset. Consequently, some branches may exhibit weak or ambiguous read support. A branch is considered to have a unique read-supported selection only when the highest-scoring candidate has positive read support and strictly exceeds the second-highest candidate support. Branches with tied or nearly indistinguishable support are therefore excluded from the comparable set, as the raw reads do not provide sufficient evidence to identify a unique preferred continuation. Among the 20,000 sampled branches, 15,319 branches in the HIV-1 dataset and 19,882 in the SARS-CoV-2 dataset yielded a unique read-supported choice, corresponding to 76.6% and 99.4% of the sampled branches, respectively. RASTRO-*S* was evaluated using context lengths (*n*) ranging from 5 to 10. In the HIV-1 dataset, match increased from 54.1% at *n* = 5 to 75.8% at *n* = 10. Similarly, in the SARS-CoV-2 dataset, match increased from 58.6% to 76.4% over the same range. These results indicate that increasing the context length improves RASTRO-*S*’s ability to distinguish between competing branch extensions.

RASTRO-*L* was evaluated under the same range of context lengths. In the HIV-1 dataset, match increased from 51.6% at *n* = 5 to 59.1% at *n* = 10, while in the SARS-CoV-2 dataset it increased from 44.87% to 49.25%. Thus, RASTRO-*L* also shows improved match with increasing context length, although the improvement is less pronounced than for RASTRO-*S*.

Overall, both models achieved match above the random-selection baseline, with performance generally improving as more sequence context was provided, Table 2 and Figure 3. This trend suggests that local sequence context provides useful information for distinguishing competing extensions at branching points. However, overall match alone is insufficient to characterize model behavior, as the branches in the evaluation set do not have equal levels of read-based evidence. For each extension option, a read score is calculated as the sum of the frequencies of its boundary *k*-mers in the corresponding raw reads. The option with the highest score is then designated as the read choice. In addition to the selected option, two quantities characterize the strength of the read-based decision: the score of the winning option and that of the strongest competing option.

**Table 2.**
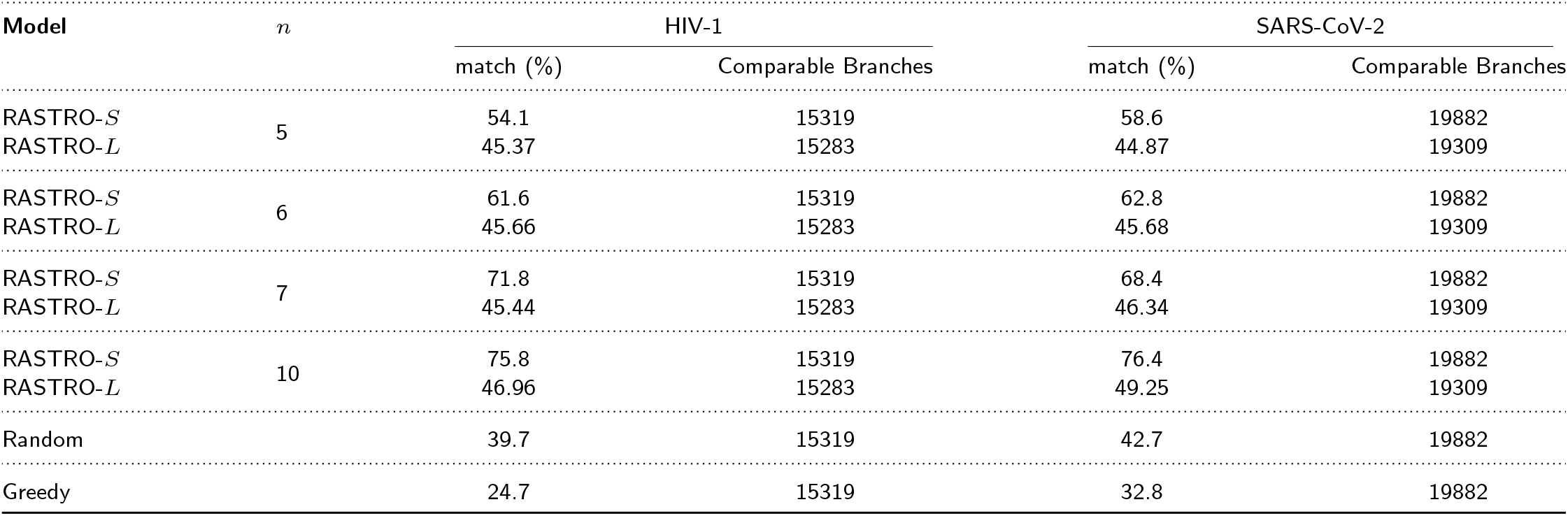
Comparison of model match with the corresponding background sequences.

| Model | $n$ | HIV-1 | | SARS-CoV-2 | |
| --- | --- | --- | --- | --- | --- |
|  |  | match (%) | Comparable Branches | match (%) | Comparable Branches |
| RASTRO-$S$ | 5 | 54.1 | 15319 | 58.6 | 19882 |
| RASTRO-$L$ | | 45.37 | 15283 | 44.87 | 19309 |
| RASTRO-$S$ | 6 | 61.6 | 15319 | 62.8 | 19882 |
| RASTRO-$L$ | | 45.66 | 15283 | 45.68 | 19309 |
| RASTRO-$S$ | 7 | 71.8 | 15319 | 68.4 | 19882 |
| RASTRO-$L$ | | 45.44 | 15283 | 46.34 | 19309 |
| RASTRO-$S$ | 10 | 75.8 | 15319 | 76.4 | 19882 |
| RASTRO-$L$ | | 46.96 | 15283 | 49.25 | 19309 |
| Random |  | 39.7 | 15319 | 42.7 | 19882 |
| Greedy |  | 24.7 | 15319 | 32.8 | 19882 |

**Figure 3.**
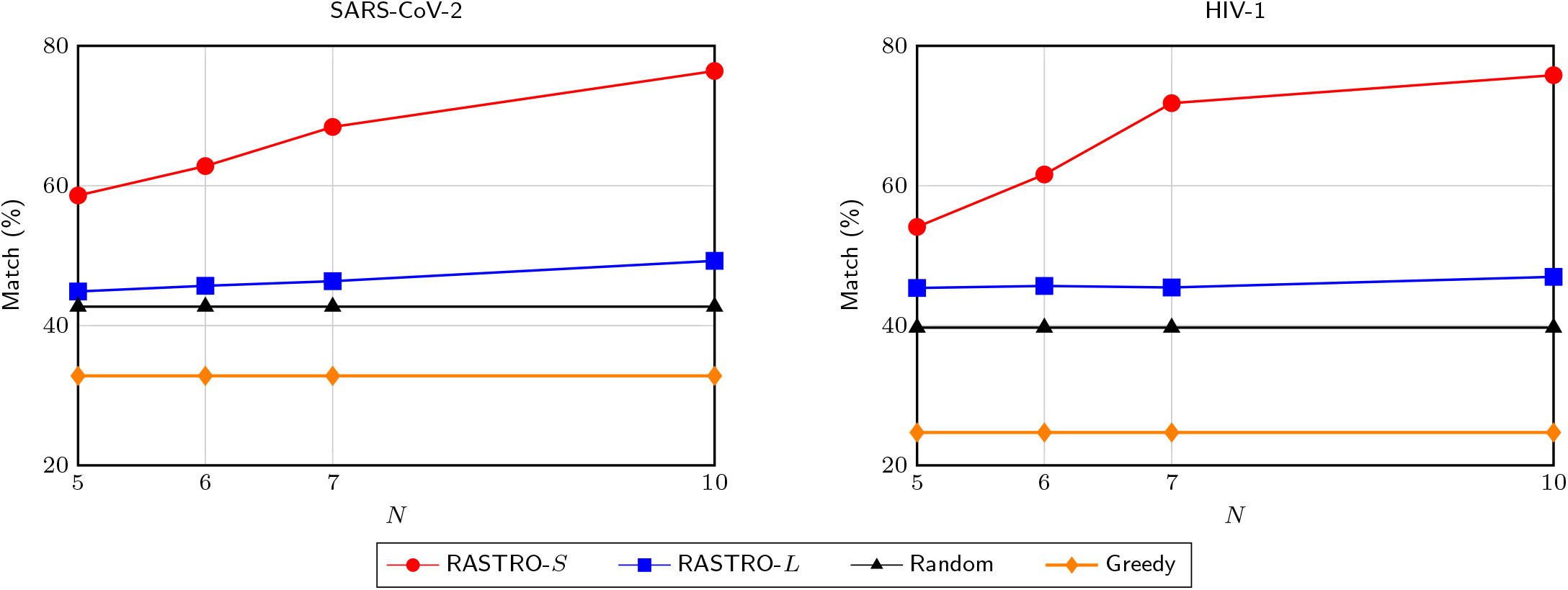
Trend of model match with increasing context length.

The difference between these two scores indicates the strength of the read evidence. When the scores are close, the read choice, although unique under our selection criterion, may be sensitive to statistical variation. Conversely, a large difference indicates stronger evidence in favor of the selected extension and therefore a more stable read-based decision. Accordingly, analyzing model match together with the strength of the underlying read evidence provides a more informative assessment of branch-level decision making.

To quantify the strength of read-based evidence, for each branch *e*, the read-support scores of the candidates are denoted by *R*_*e*_, where *e* represents a candidate extension. The largest and second-largest read scores are defined as 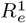 and 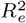, respectively. The evidence strength ratio, referred to as margin_ratio, is then calculated:

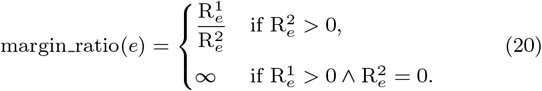

A larger value indicates a stronger preference of the reads toward the selected candidate. Thresholding is applied by selecting a fixed value *T* (for example, *T* ∈ {2, 5, 10}) and calculating the match only over branches satisfying margin_ratio(*e*) ≥ *T*.

The set of comparable branches is defined as:

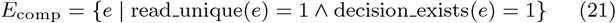

By applying a threshold *T*, the subset of branches with stronger read evidence is obtained as:

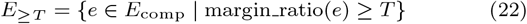

Finally, the thresholded match for a given threshold *T* is calculated using the same definition as the overall match, with the numerator and denominator restricted to the branches in *E*_≥*T*_:

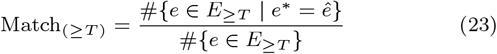

A threshold *T* is applied to the margin ratio to identify branches with stronger read evidence:

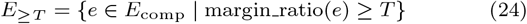

Thresholded match is then computed using the same definition as overall match, but restricted to *E*_≥*T*_:

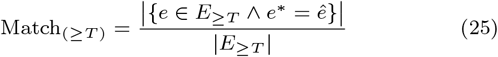

Thus, increasing *T* progressively restricts the evaluation to branches for which the read evidence provides a more distinct preference between competing extensions. This thresholding provides a robustness analysis of model behavior under varying levels of read-based evidence, Table 3 and Figure 4. For both RASTRO-*S* and RASTRO-*L*, match increases monotonically as *T* increases, indicating that both models perform better when the underlying read evidence provides a clearer distinction between competing paths. In contrast, the random baseline does not exhibit a systematic increase across these subsets and remains substantially lower. The greedy baseline also remains below the random baseline, indicating that selecting the longest candidate extension alone provides less useful information than chance-level selection for identifying read-supported branch decisions.

**Table 3.** Analysis of model match based on different *margin_ratio* thresholds across different context lengths.

| Model | HIV-1 |  | SARS-CoV-2 |  |
| --- | --- | --- | --- | --- |
| | *margin_ratio* $\geq 5$ | *margin_ratio* $\geq 10$ | *margin_ratio* $\geq 5$ | *margin_ratio* $\geq 10$ |
| *n* = 5 |  |  |  |  |
| RASTRO-*S* | 54.5 | 54.8 | 62.4 | 63.8 |
| RASTRO-*L* | 45.81 | 46.33 | 41.16 | 42.67 |
| *n* = 6 |  |  |  |  |
| RASTRO-*S* | 68.2 | 70.7 | 68.0 | 70.1 |
| RASTRO-*L* | 45.53 | 45.98 | 42.10 | 43.65 |
| *n* = 7 |  |  |  |  |
| RASTRO-*S* | 76.6 | 81.1 | 74.9 | 77.6 |
| RASTRO-*L* | 46.08 | 46.27 | 43.02 | 44.60 |
| *n* = 10 |  |  |  |  |
| RASTRO-*S* | 89.8 | 94.4 | 84.9 | 87.9 |
| RASTRO-*L* | 48.25 | 48.34 | 48.26 | 49.14 |
| Random | 41.1 | 41.5 | 41 | 40.3 |
| Greedy | 14.6 | 11.6 | 25.9 | 22.3 |

**Figure 4.**
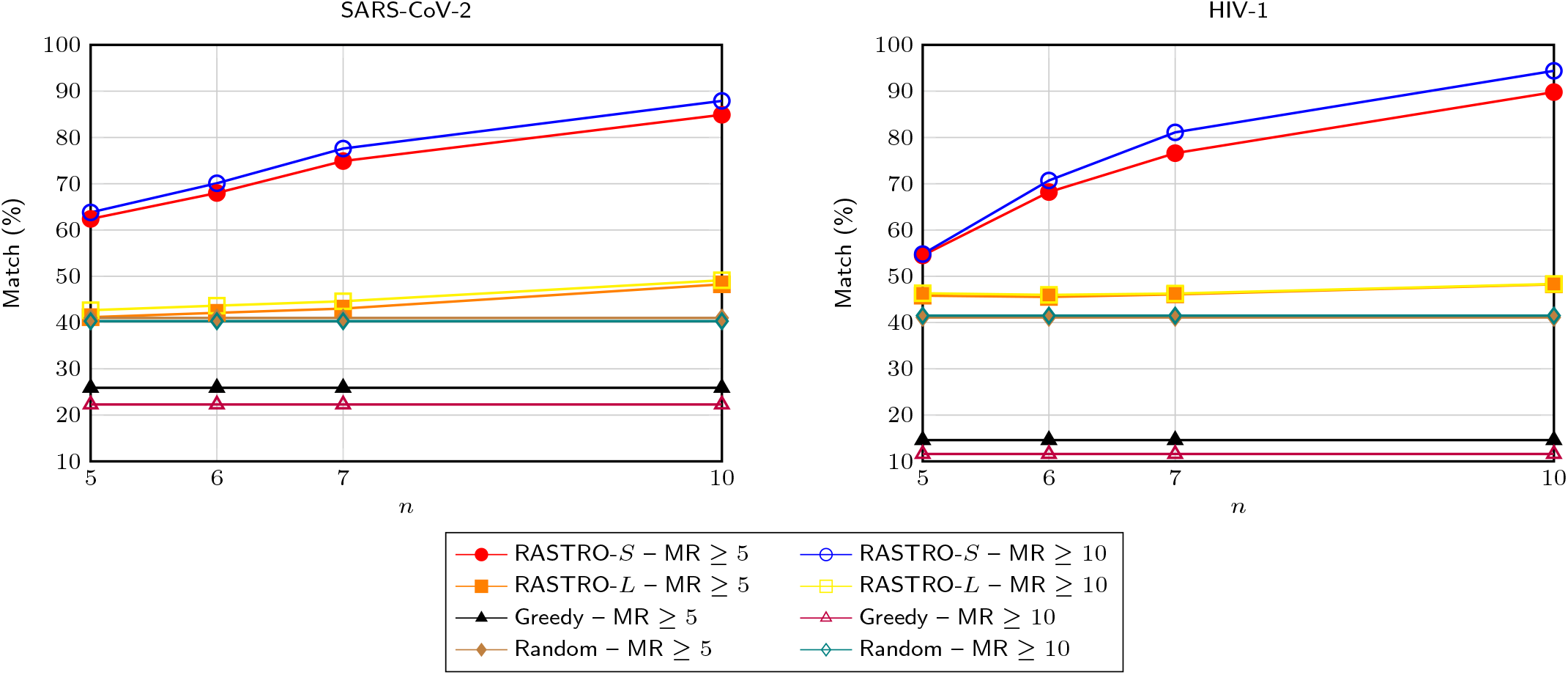
Change in model match with increasing context length under different *margin_ratio* thresholds.

## Conclusion

The performance of RASTRO-*S* was evaluated by comparing its branching decisions with read-supported evidence. RASTRO-*S* consistently outperformed the random baseline across the evaluated context lengths and datasets and achieved higher match with read evidence than RASTRO-*L* in most settings. Increasing the context length from *n* = 5 to *n* = 10 generally improved RASTRO-*S*’s match in both datasets, indicating that longer sequence contexts provide additional information for branching-point decisions. Moreover, increasing the *margin_ratio* threshold from ≥ 5 to ≥ 10 consistently increased RASTRO-*S*’s match, demonstrating that stronger read evidence is associated with more reliable model decisions.

The observed match should not be expected to reach 100%, since model-based scores and read-supported choices capture different aspects of the sequencing data. Unitigs represent a compressed representation of local *k*-mer connectivity, whereas the read-supported choice is explicitly frequency-based and determined from the occurrence of boundary *k*-mers in the raw reads. Consequently, the two criteria may differ at branches with multiple plausible paths, heterogeneous coverage, polymorphic signals, or sampling effects. To ensure a reliable evaluation, only branching points with a unique and unambiguous read-supported choice were included.

The higher match of RASTRO-*S* compared with RASTRO-*L* can be explained by their different modeling strategies. RASTRO-*S* is trained on data from the same sample and therefore captures sample-specific sequence patterns, whereas RASTRO-*L* is a pretrained model that provides broader sequence-level priors. Thus, in the setting evaluated here, the lightweight sample-centric RASTRO-*S* scorer can align more closely with sample-specific read evidence than a larger general-purpose model. This result should not be interpreted as general superiority over larger language models, but rather as evidence for the value of sample-centric modeling in branching-point decision-making.

Raw reads were used as the evaluation evidence rather than final contigs because branching points are precisely where competing paths exist and where subsequent assembly decisions may conceal underlying ambiguity. Evaluating against contigs could therefore measure consistency with a particular assembly or alignment output rather than with the underlying sequencing evidence. In contrast, the proposed read-choice criterion uses boundary *k*-mer occurrences to quantify support for each candidate continuation, providing a reproducible and lightweight evaluation that remains closely tied to the raw data.

Several limitations should be considered. The computational cost of some comparative experiments restricted the number of replicates, the size of the evaluated genomes, and the range of methods that could be tested. Dataset availability also limited the evaluation, as suitable datasets needed to provide compatible raw reads and unitig-graph information while satisfying requirements related to read type, format, and preprocessing. Furthermore, although RASTRO-*S* was evaluated on both HIV-1 and SARS-CoV-2 and showed similar performance trends relative to the random baseline, two datasets are insufficient to establish broad generalizability. Evaluation on a larger and more diverse collection of genomes and sequencing conditions is therefore needed.

Attempts were also made to compare RASTRO-*S* with tools such as ALE, SAMA, GABI, and SPAdes. However, these approaches address different levels of the assembly workflow, primarily producing or evaluating contigs or complete assemblies rather than making decisions at unitig graph branching points. Consequently, direct comparison of their outputs with RASTRO’s branching decisions would not constitute a methodologically equivalent evaluation.

Although RASTRO is not currently a complete genome assembler, its branching-point scores and read-evidence consistency measures provide a foundation for integrating sequence-based scoring into broader assembly workflows. This provides a practical direction for combining RASTRO with existing graph-processing and assembly workflows, where RASTRO can serve as an additional decision signal alongside coverage, paired-end information, graph topology, and other sources of evidence. To test this and further analyze the unitig-graph datasets and provide a consistent representation for downstream analysis, we independently used agtools (Mallawaarachchi et al., 2026), a general-purpose framework for manipulating assembly graphs generated by different assemblers. Agtools provides functionality for loading, traversing, and processing assembly-graph representations, including unitig graphs, thereby allowing the graph structure and connectivity of the datasets to be accessed in a standardized manner. The resulting graph representation was used to identify branching points and construct candidate extensions, which were subsequently scored by RASTRO-*S* and RASTRO-*L*. Future work should investigate this integration at larger scale and evaluate whether RASTRO scores can improve path ranking and branch resolution within complete assembly pipelines. Further evaluation across a broader range of genomes, sequencing conditions, and genomes not represented in the training data will also be necessary to assess the generalizability of the approach.

## Code Availability

The source code is available at https://github.com/BioinformaticsIASBS/RASTRO.

## Data availability

The sequencing datasets used in this study are publicly available under the NCBI Sequence Read Archive (SRA) accession numbers `SRR7693968` and `SRR10605873`. The corresponding unitig graphs are available from Logan (https://logan-pub.s3.amazonaws.com/), and the raw sequencing reads can be accessed from the NCBI SRA and the European Nucleotide Archive (ENA (https://www.ebi.ac.uk/ena/).

## Author contributions statement

M.H. conceived and defined the research project and supervised the study. S.B. and M.H. contributed to the conceptualization of the methodology. S.B. carried out the implementation and experimental analysis. Both authors contributed to the writing, review, and editing of the manuscript.

